# Proteomic characterization of sporadic clear cell renal cell carcinoma reveals a matrix-dense pseudocapsule and heterogeneous tumor subtypes

**DOI:** 10.1101/2025.10.23.683844

**Authors:** Tobias Feilen, Grigor Andreev, Manuel Rogg, Miguel Cosenza-Contreras, Niko Pinter, Max Zirngibl, Antonia Elsässer, Pitter Huesgen, Markus Grabbert, Christoph Schell, Oliver Schilling

## Abstract

Clear cell renal cell carcinoma (ccRCC) frequently develops a fibrotic pseudocapsule (PC) at the tumor boundary, yet its implications in the disease remain incompletely defined. We performed DIA-based proteomics on formalin-fixed, paraffin-embedded (FFPE) specimens of sporadic ccRCC from 153 patients, comprising 142 tumor, 123 pseudocapsule (PC), and 121 non-malignant adjacent tissue (NAT) samples.

The PC displayed a distinctive proteomic signature characterized by matrix accumulation, enhanced remodeling and ECM-cell signaling, consistent with a signaling-competent boundary and active growth factor sequestration rather than a passive fibrotic barrier. Within the tumor, we observed canonical metabolic reprogramming with upregulated glycolysis and hypoxia markers and suppression of aerobic metabolism, accompanied by strong cell-cycle and pro-angiogenic signatures. Immune programs included increased antigen processing and presentation, together with elevated inflammasome and pyroptosis signatures. T-cell markers were enriched in the PC, indicating an immune-active boundary. Among tumors, we identified five distinct proteomic subtypes (C1–C5) spanning proliferative/mitotic, metabolic, immune/EMT/mTOR, and ECM phenotypes. Semi-specific peptide analysis indicated elevated endogenous proteolysis in the tumor and varied across clusters, with C1 and C5 showing the greatest proteolytic burden. Mechanotransduction features showed only modest PC elevation but were dominated by inter-patient variation.

The present study defines a matrix-rich, signaling-active pseudocapsule and a classification of heterogeneous tumor subtypes in sporadic ccRCC. The framework provides a compartment-resolved, cluster-informed approach, and it highlights actionable axes of potential clinical relevance.

## Introduction

Renal cell carcinoma (RCC) ranks among the ten most common malignancies worldwide. In 2020, more than 430,000 individuals were diagnosed with RCC, with men affected approximately twice as often as women.^1^ The most prevalent histological subtype is clear cell RCC (ccRCC), which accounts for over 70% of all RCC cases.^1^ A histo-morphological hallmark feature of ccRCC, present in over 90% of cases, is the formation of a pseudocapsule (PC), a capsule-like boundary zone that separates the tumor from the non-malignant renal parenchyma.^2^ The PC forms upon the expansive tumor growth, which exerts compression forces on surrounding tissue.^3^ Histologically, this boundary region is characterized by compressed and remodeled parenchyma containing collagen fibers, smooth muscle bundles, and fibrous tissue that develops in response to ischemia and compression-induced necrosis. Clinically, the PC serves as an important surgical and pathological landmark. An intact and fibrotic PC acts as a physical barrier that can aid nephron-sparing surgery and is associated with a more localized tumor growth, whereas a disrupted or less fibrotic PC correlates with invasive tumor behavior, greater metastatic potential, and a shorter progression-free survival.^4–7^ Despite its potential clinical significance, the molecular composition of the PC in sporadic ccRCC remains insufficiently characterized. A deeper understanding of the PC’s molecular landscape will provide insight into its role as a structural and functional barrier and into how it contributes to tumor progression. At the molecular level, ccRCC is typically initiated by biallelic *VHL* inactivation. Loss of the pVHL protein prevents oxygen-dependent degradation of hypoxia-inducible transcription factors (HIF-1α and HIF-2α). Their stabilization drives a pseudo-hypoxic transcriptional program that promotes angiogenesis, cell survival, and metabolic rewiring.^8,9^ Beyond *VHL*, various driver mutations in chromatin and epigenetic regulators, including *PBRM1*, *SETD2*, *BAP1*, and *KDM5C*, contribute to tumor development and heterogeneity, with known associations to distinct biological behaviors and clinical outcomes.^10,11^ In addition, HIF signaling profoundly reshapes the matrisome, the composite of extracellular matrix (ECM), basement-membrane proteins, and ECM-associated regulators such as proteoglycans, glycoproteins, and proteases.^12–14^ Increasing evidence indicates that the tumor-associated matrisome actively modulates hallmarks of cancer, including proliferation, angiogenesis, invasion, metabolic rewiring, and immune evasion.^15^ Here, we present a compartment-resolved proteomic characterization of tumor, PC, and NAT in sporadic ccRCC. Building on our previous analysis of VHL disease-associated ccRCC, we sought to define the proteomic landscape of the PC in the much more prevalent sporadic disease.^16^

## Materials and Methods

### Ethics Statement

Ethics approval was obtained from the Ethics Committee of the University Medical Center Freiburg (576/17). All participants provided written informed consent permitting the use of their tissue specimens for research. Patient information was pseudonymized prior to study inclusion.

### Tissue Collection, Fixation, and Dissection

Tissue specimens were obtained between 2008 and 2021 during surgical resection of tumors at the University Medical Center Freiburg. Following resection, tissues were placed in formalin solution and paraffin-embedded. Tissue specimens were sectioned, processed, and analyzed following standard protocols for histopathological diagnostics. For proteomic analysis, 10 µm thick formalin-fixed paraffin-embedded (FFPE) sections were automatically deparaffinized and compartment-specific dissection was performed using a binocular microscope and under the guidance of matched hematoxylin and eosin (H&E)-stained sections by an experienced pathologist to isolate non-malignant adjacent tissue (NAT), pseudocapsule (PC) and tumor regions. A representative H&E image of the tumor, PC, and NAT is shown in **Fig. S1**.

### Sample Preparation for LC-MS/MS Analysis and Data Acquisition

Following dissection, tissue specimens were transferred into lysis buffer (100 mM HEPES, 1% (w/v) SDS, pH 8). Heat-induced antigen retrieval (HIAR, 2 h. 95 °C) was performed, followed by ultrasonication in a Bioruptor Plus (Diagenode, Belgium; 20 cycles, 30 s on, 30 s off). Protein concentrations were determined by the BCA assay (Thermo Fisher Scientific, Germany), yielding 50–150 µg of protein per sample, of which 25 µg in 150 µL lysis buffer were further subjected to automated processing using the automated liquid handling platform AssayMAP Bravo (Agilent Technologies, USA). Proteins were reduced with 5 mM TCEP and 20 mM chloroacetamide (CAA) for 30 min at 37 °C in the dark. Protein binding and cleanup were done using the SP3 protocol as published previously.^17^ In brief, Sera-Mag magnetic carboxylate beads were added at 3.1 µg/µL, following ACN addition to 60% (v/v) final concentration and shaking at 1050 rpm for 10 min. Beads were washed three times (2x EtOH 70%, 1x ACN 100%) and resuspended in 100 µL ammonium bicarbonate (ABC) buffer (100 mM, pH8). Proteolysis proceeded with lysyl endopeptidase (Lys-C, Wako Chemicals, Germany) for 2 h at 42 °C and 800 rpm, followed by trypsin (Serva, Germany) for 15 h at 37 °C and 800 rpm at a 1:100 and 1:25 enzyme-to-protein ratio, respectively. Beads were separated from the supernatant using a magnetic rack, and peptide concentration was determined by the peptide BCA assay (Thermo Fisher Scientific, Germany). 500 ng of peptides were spiked with 100 fmol of indexed retention time (iRT) peptides and loaded onto Evotips (Evosep, Denmark) following the manufacturer’s protocol.

### LC-MS/MS Measurement

LC-MS/MS analyses were carried out on a TimsTOF Flex Mass Spectrometer (Bruker Daltonics, Bremen, Germany) equipped with a CaptiveSpray ion source. Peptides were separated on an Evosep One HPLC system using the 30SPD method (44 min gradient, 500 nL/min) coupled to an EV1137 performance 15 cm reversed-phase column (Evosep). Mobile phases were: 0.1% aqueous formic acid (buffer A) and 0.1% formic acid in can (buffer B). Method optimization included optimal placement of DIA-PASEF isolation windows prior to the measurements using the py_diAID algorithm.^18^ The mass-spectrometer was operated in positive dia-PASEF mode with a 100–1700 *m*/*z range* for MS1, 400–1200 *m*/*z range* for MS2 scans, and 0.7–1.3 V*s/cm^2^ ion mobility (1/K0) range. Collision-induced dissociation energy was set to 20-59 eV, ion accumulation time to 100 ms with a 100% duty cycle. The capillary voltage was set to 1600 V, and the drying gas flow rate was 3 l/min (180 °C drying temperature). For DIA measurements, 50 mass windows with optimized *m*/*z* range were applied, giving a total cycle time of 2.76 s.

### Data Analysis

Raw LC-MS/MS data were processed with DIA-NN (v1.9.2).^19,20^ A predicted spectral library was generated from the human reference proteome database (downloaded from https://www.ebi.ac.uk/reference_proteomes/ on 22/07/2023), supplemented with common contaminants and 11 iRT peptides. The match between runs (MBR) algorithm was enabled. For peptide precursors, a range of 7–30 amino acids and charge states of 1–4 were included, with an *m*/*z* window of 100–1700, allowing one missed cleavage. Fragment ion *m*/*z* ranged from 390–1210, and identifications were filtered at a false discovery (FDR) threshold of 1%. Statistical and exploratory analyses were conducted in RStudio (v4.3.0) using in-house scripts and publicly available R packages. Quantitative proteomic data were extracted using the DIA-NN R package (v1.0.1), applying the MaxLFQ algorithm to estimate protein abundances from proteotypic peptides only.^19,21^ Protein intensities were log_2_-transformed prior to analysis. The mixOmics package (v6.24.0) was utilized for supervised and unsupervised exploratory statistics.^22^ Differential abundance analysis was performed with limma (v3.56.2), and gene set enrichment analysis was carried out using clusterProfiler (v4.8.3), incorporating Gene Ontology, hallmark and curated cancer cell atlas (3CA) databases (MSigDB collections).^23–27^ Missing values were imputed using MissForest (v1.5).^28^ To correct batch effects and to align datasets, HarmonizR (v1.0.0) was applied, and ECM proteins were classified via MatrisomeAnalyzeR (v1.0.1).^29^ Monte Carlo reference-based consensus clustering of tumor samples was performed with M3C (v1.22.0), and variance partitioning analysis was conducted using the variancePartition R package (v1.32.5).^30,31^

## Results and Discussion

### Sporadic ccRCC Patient Cohort and Proteome Coverage

For this study, we assembled a sporadic ccRCC cohort based on archived patient material obtained from the Department of Pathology at the University Medical Center Freiburg, including tumor, pseudocapsule (PC), and non-malignant adjacent tissue (NAT) samples, to enable a compartment-resolved proteome profiling (**Fig. 1A**). The cohort initially comprised 154 ccRCC patients. As some compartments were unavailable and post-analysis quality control excluded technical outliers and low-quality samples, the final dataset included 386 FFPE specimens from 153 patients, (142 tumor, 123 PC, and 121 NAT samples). The cohort included 95 males and 58 females with a median age of 63 years (IQR interquartile range = 57–70) and 66 years (IQR = 57–76), respectively. Further cohort information can be found in **Table S1**.

**Figure 1.**
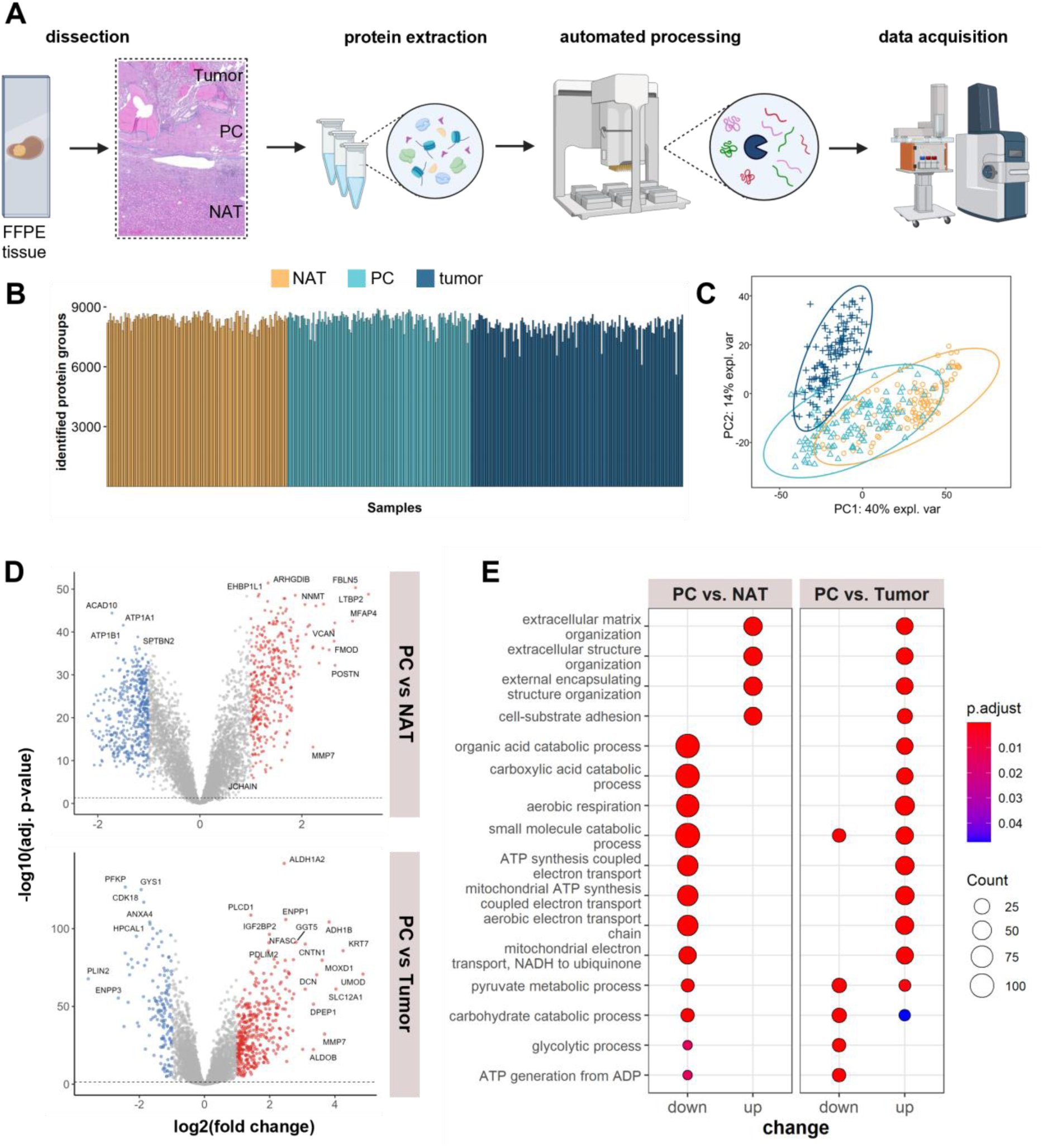
Sporadic ccRCC cohort: **A** Overview of experimental workflow, including tissue dissection, separating NAT, PC, and tumor, protein extraction, automated sample processing with tryptic digestion, followed by MS measurement and data analysis. **B** Identified protein groups across all samples of the cohort. **C** Principal component analysis (PCA) of tumor (dark blue), PC (light blue), and NAT (orange) samples. Ovals indicate 95% confidence intervals. **D** Volcano plots showing differentially regulated proteins between PC and NAT (top) and PC and tumor (bottom). Proteins on the right are upregulated in the PC (fold change ≥ |2|, BH adj. p-value ≤ 0.05). E GO enrichment analysis of upregulated and downregulated biological processes in PC vs NAT and PC vs tumor (BH adj. p-value ≤ 0.05). Created with Biorender.com.

Across all specimens, we identified 9857 protein groups at a 1% false discovery rate (FDR), with a per-sample median of 8311 (IQR = 7841–8781; **Fig. 1B**). Principal component analysis (PCA) revealed that tumor and NAT samples formed discrete clusters, whereas PC profiles localized between and within them with a strong overlap to NAT (**Fig. 1C**). To balance depth and completeness for downstream analyses, we compiled two matrices: First, a stringent set requiring > 70% completeness within each compartment for proteins, yielding 7167 quantified proteins suitable for robust differential analyses. Second, an extended set retaining proteins present in > 50% of samples in any one compartment to support broader pathway-level discovery, yielding 8919 proteins after imputation. This two-tier strategy accommodates tissue-specific expression patterns while obviating bias from missingness.

To contextualize our findings, we integrated a publicly available CPTAC ccRCC proteome resource with our previously published syndromic ccRCC dataset and the present sporadic ccRCC cohort using ComBat-based batch correction.^16,32,33^ PCA showed substantial overlap of tumor and NAT samples across studies (**Fig. S2**), indicating that the proteomic features observed in our data are representative of ccRCC biology and further supporting the robustness of our study’s findings.

### The Proteomic Signature of the Pseudocapsule

To further deepen our understanding of the significance of the PC in sporadic ccRCC, we performed in-depth proteomic analysis of 123 PC specimens. To define PC-specific biology in sporadic ccRCC, we compared PC to NAT and tumor. Both contrasts indicated a PC proteome that is vastly different from tumor and NAT (**Fig. 1D**). Gene ontology (GO) enrichment analysis highlighted terms linked to extracellular matrix (ECM) organization, external encapsulating structure organization, and cell-substrate adhesion among the most significantly increased biological processes (**Fig. 1E**, left). Conversely, proteins assigned to mitochondrial and aerobic energy metabolism, including aerobic respiration, mitochondrial electron transport, and ATP synthesis coupled to electron transport, were depleted in PC vs NAT (**Fig. 1E**, left).

Against the tumor, the PC showed lower abundance of pathways indicative of glycolytic metabolism and carbohydrate catabolism, whereas several oxidative phosphorylation-associated processes were elevated (**Fig. 1E**, right). Together with the NAT contrast, these results position the PC as an intermediate but distinct compartment. It is enriched in ECM and cell-adhesion components and demonstrates suppression of mitochondrial programs vs NAT. The depletion of glycolytic pathways vs tumor potentially reflects the strong canonical shift toward anaerobic metabolism in ccRCC tumors. This PC pattern mirrors the proteomic profile observed in our previously published cohort of syndromic ccRCC, where the PC similarly combined a fibrotic and adhesive program with reduced mitochondrial functions.^16^

### Proteins Involved in ECM Remodeling and Fibrotic Signaling Characterize the PC Proteome

To isolate a PC-specific signature, we intersected proteins significantly increased in the PC relative to both tumor and NAT. This yielded 308 PC-enriched proteins, of which one third map to the matrisome (**Fig. 2A, S3**). We observed a broad PC-specific elevation of ECM scaffolding proteins, most prominently collagens. Notably, the collagen chains COL6A1/2/3 are PC enriched, consistent with prior reports of the tumor boundary.^34^ Beyond collagens, fibronectin (FN1), elastin (ELN), fibrillin-1 (FBN1), and multiple laminins show concordant enrichment in the PC. Small leucine-rich proteoglycans (SLRPs; e.g., DCN, LUM, BGN, FMOD, PRELP) are likewise elevated, supporting roles in fibrillogenesis, fibril stabilization, and matrix organization within the boundary zone.^35^ We next examined enzymes orchestrating ECM synthesis, remodeling, and signaling (**Fig. 2B**, left). Surprisingly, enzymes involved in collagen maturation, including prolyl-4-hydroxylases (P4HA1/2 and P4HB), lysyl hydroxylases (PLOD1/2/3), the lysyl oxidase (LOX), the collagen chaperone FKPB10, and the crosslinking enzyme TGM2, are more abundant in the tumor compared to the PC (**Fig. 2A**). Within the remodeling axis, both the PC and the tumor exhibit high overall abundance of matrix metalloproteinases (MMPs), albeit with distinct profiles. Several MMPs are enriched in both compartments, whereas MMP7 and MMP23B exhibit inverse regulation (upregulated in PC, downregulated in tumor). Consistent with these observations, semi-specific peptide analysis identifies a higher number of semi-specific (i.e., proteolytically truncated) collagen peptides in the PC **(Fig. 2C**). This suggests that the PC-enriched tissue inhibitors of metalloproteinases (TIMPs 1–3; **Fig. 2B**, middle) do not lead to complete restriction of collagen proteolysis at the boundary and indicate spatially regulated proteolysis. Building on these remodeling dynamics, we next examined ECM-associated signaling components. The PC is characterized by an elevated abundance of multiple integrin subunits, syndecans, and CD44, as well as matrix-tethering growth factor modulators including latent TGFβ-binding proteins (LTBPs) and glypicans (**Fig. 2B**, right). By shaping tumor growth factor bioavailability and receptor interactions, these factors likely contribute to a distinct microenvironment at the tumor margin that remains highly responsive to extracellular cues.^36,37^

**Figure 2.**
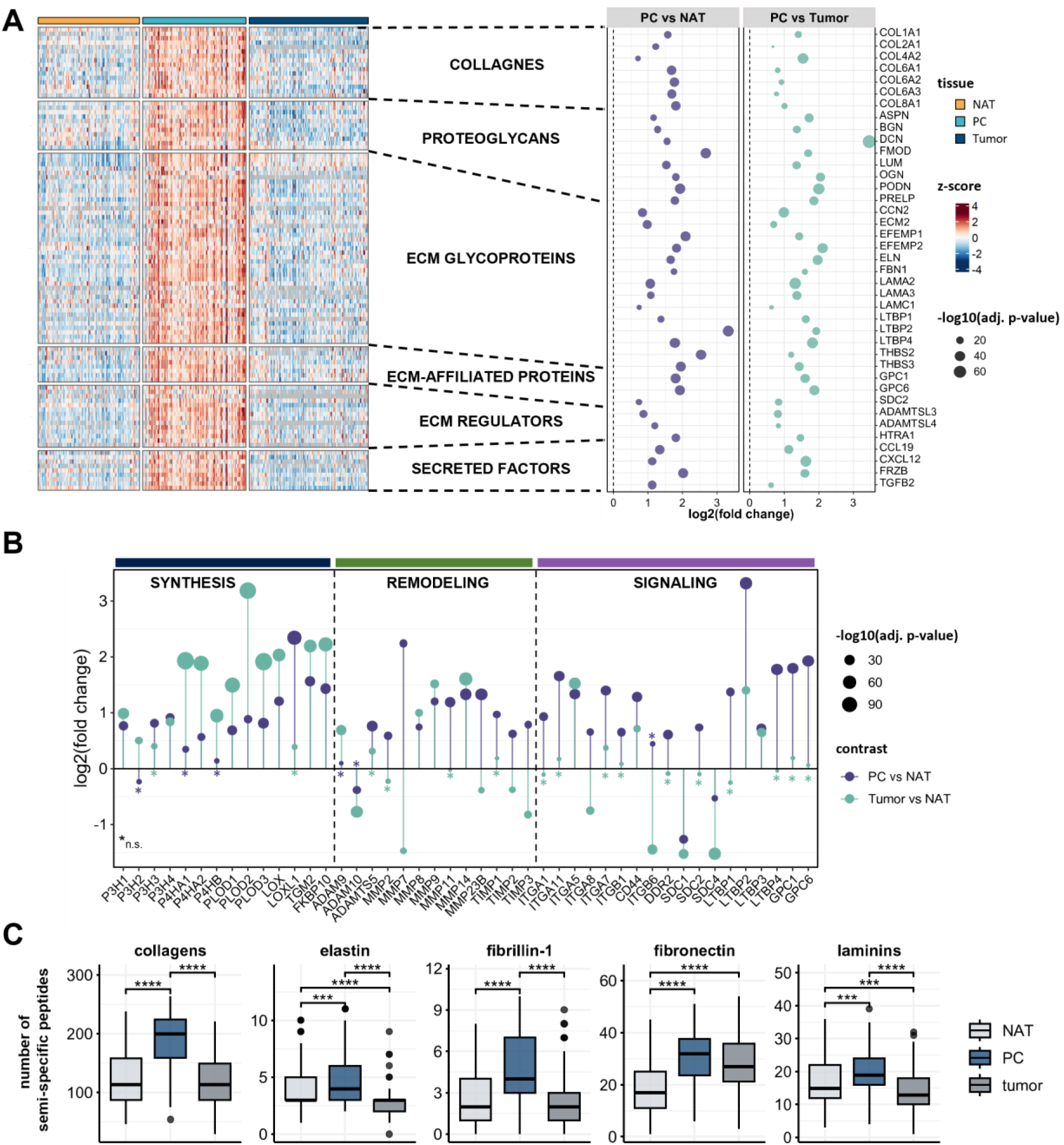
The matrisomal landscape of the PC: **A** Heatmap (left) comparing the abundance of PC enriched matrisomal proteins and a dotplot (right) showing the log2 fold changes of selected matrisomal proteins in PC and tumor vs NAT. **B** Dotplot of selected differentially regulated proteins involved in ECM synthesis (left), remodeling (middle) and signaling (right) in PC vs NAT (blue) and tumor vs NAT (turquoise; imputed dataset, fold change ≥ |1.5|, BH adj. p-value ≤ 0.05, asterisks indicate non-significant changes, colors indicate the respective contrast). **C** Boxplots showing the number of identified semi-specific peptides of selected ECM scaffolding proteins; **p < 0.01, ***p < 0.001; ****p < 0.0001 (Dunn’s test). Created with Biorender.com.

Further highlighting the specialized nature of the PC, its proteome is enriched in canonical fibrotic drivers, including TGFβ2 and CCN2 together with fibroblast-associated markers (S100A4/FSP1, PDGFRB, and CD248) and a panel of cancer-associated fibroblast (CAF) subtype proteins (**Fig. 3A**). Stratification by CAF class showed a dominant myofibroblast-like CAF (myoCAF) signature, e.g., POSTN, FAP, FN1, and collagens (COL1A1, COL3A1), whereas inflammatory CAF (iCAF) markers were only modestly increased (C3, CXCL12) or reduced (DPP4, CFD). These patterns are consistent with a contractile, matrix-depositing stroma at the tumor boundary that helps to establish the PC’s matrix-dense niche. Such myoCAF enrichment may increase local stiffness and promote growth factor sequestration and presentation (e.g., TGFβ), thereby creating a signaling-rich interface rather than a purely structural capsule.

**Figure 3.**
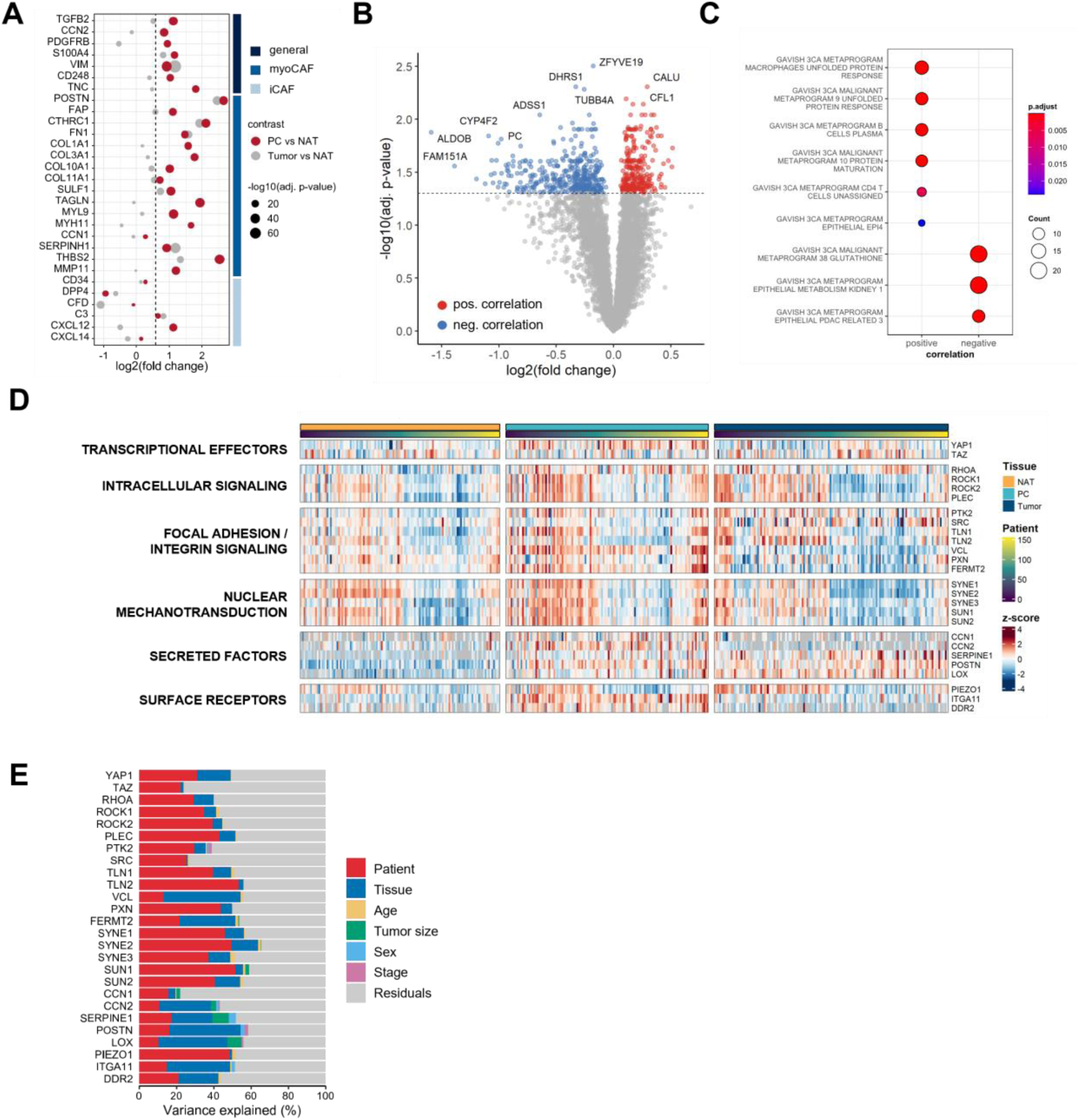
Tumor size correlation analysis and mechanotransduction: **A** Dotplot of selected significantly upregulated general fibrosis-associated proteins, markers for myofibroblast-like cancer-associated fibroblasts (myoCAF) and inflammatory CAFs (iCAF) in the PC vs NAT (imputed dataset, fold change ≥ 1.5, BH adj. p-value ≤ 0.05). **B** Volcano plot showing PC proteins positively (red) and negatively (blue) correlated with tumor size (BH adj. p-value ≤ 0.05). **C** Enrichment analysis of Hallmark gene sets (MSigDB) among PC proteins exhibiting positive or negative correlation with tumor size. **D** Heatmap comparing the abundance of selected mechanotransduction proteins in tumor, PC, and NAT. **E** Variance partitioning analysis showing the proportion of expression variance in mechanotransduction-associated proteins.

### Tumor Size-Dependent Alterations of the PC Proteome and Mechanotransduction Signatures

To assess whether tumor expansion reshapes the PC, we correlated tumor diameter with the PC protein abundance (**Fig. 3B**). We detected 271 proteins that increased and 410 that decreased with tumor size (BH adj. p-value ≤ 0.05). Proteins positively associated with tumor size were enriched for unfolded protein response, indicative of cellular stress, and immune programs involving B cells and CD4 T cells (**Fig. 3C**). Conversely, negatively associated sets included glutathione metabolism components and epithelial kidney metabolic programs. Reduced representation of the glutathione pathway may indicate diminished antioxidant buffering in the mechanically compressed boundary, potentially increasing oxidative stress sensitivity. Decreased epithelial metabolic programs likely reflect loss of differentiated nephron features in the PC as compression and ischemia progress, consistent with fibrotic and inflammatory remodeling. Together, these size-dependent trends suggest that mechanical stress imposed by tumor growth actively modulates the boundary proteome.

To further explore the effect of mechanical stress on the PC proteome, we profiled proteins linked to different categories of mechanotransduction (**Fig. 3D**). While the transcriptional effectors YAP1 and TAZ did not show a PC-specific enrichment, we observed a modest elevation of intracellular signaling (e.g., ROCK1/2, PLEC), focal adhesion/integrin signaling (e.g., TLN1/2, VCL, FERMT2), and nuclear mechanotransduction (e.g., SYN1/2/3, SUN1/2) proteins in the PC. Notably, the patient-wise sorted heatmap revealed coordinated elevations across all three tissues within certain individuals. We quantified these observations using variance partitioning analysis with a linear mixed model including patient, tissue, tumor size, age at diagnosis, sex and tumor stage (**Fig. 3E**). Across many markers, including YAP1, RHOA, ROCK1/2, PTK2, TLN1/2, PXN, SYN1/2/3, SUN1/2, and PIEZO1, the dominant source of explained variance was the patient, whereas the tissue type contributed relatively little. These results indicate that inter-patient variability, potentially reflecting differences in stromal composition and tissue mechanics, exerts a stronger impact on mechanotransduction signatures than local tissue context.

### Proteome Profiling Reveals Core Metabolic, Proliferative, Angiogenic, and Immune Hallmarks of ccRCC

Differential abundance analysis of tumor versus NAT highlighted the metabolic hallmarks of sporadic ccRCC (**Fig. 4A**). Key glycolytic enzymes and hypoxia-associated markers were elevated, whereas tricarboxylic acid (TCA) cycle and electron transport chain proteins were broadly reduced. Together, these contrasts recapitulate the pseudo-hypoxic and glycolytic signatures of ccRCC and provide a proteomic corroboration for the subsequent analyses.

**Figure 4.**
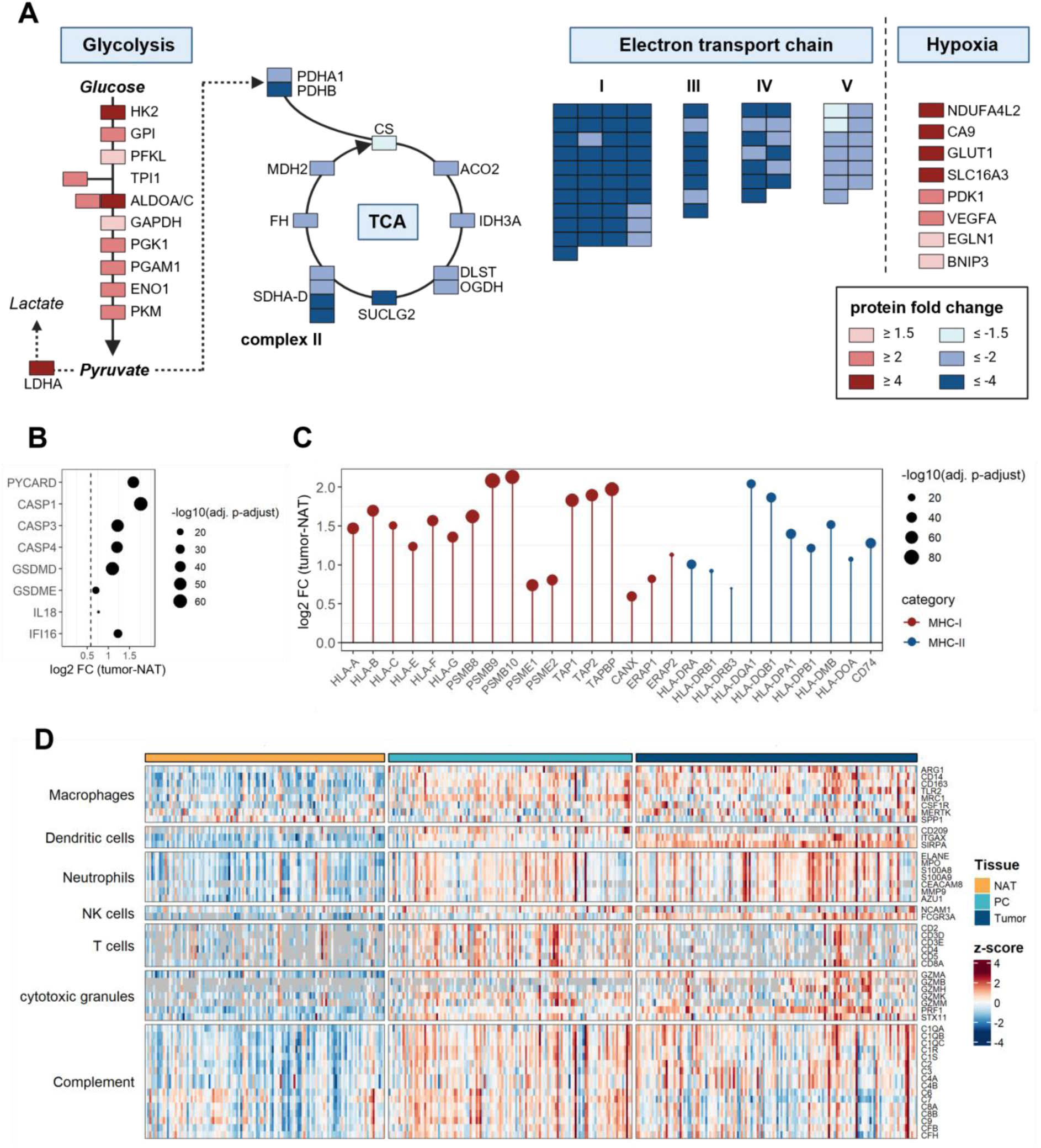
The ccRCC tumor landscape: **A** Schema showing the protein log2 fold changes between tumors and NAT of selected metabolic pathways (glycolysis, tricarboxylic acid (TCA) cycle, and electron transport chain) and hypoxia markers (BH adj. p-value ≤ 0.05). **B** Dotplot of significantly upregulated inflammasome and pyroptosis-associated proteins in the tumor (fold change (FC) ≥ 1.5, BH adj. p-value ≤ 0.05). **C** Dotplot of significantly upregulated proteins involved in antigen-processing and presentation in the tumor (imputed dataset, fold change (FC) ≥ 1.5, BH adj. p-value ≤ 0.05). **D** Heatmap comparing the abundance of proteins associated with the presence of immune cells, and complement activity in tumor, PC, and NAT. Created with Biorender.com.

We noted that tumor tissue displayed a higher abundance of proteins central to inflammasome formation and pore-forming cell death (**Fig. 4B**). The adaptor PYCARD and the DNA sensor IFI16 were increased, alongside CASP1/4 and gasdermin D, indicating capacity for both canonical and non-canonical pyroptosis.^38^ In parallel, CASP3 and gasdermin E were elevated, compatible with an apoptosis-pyroptosis switch in which caspase-3 cleavage of gasdermin E can trigger secondary pyroptosis.^39^ These data suggest that tumor regions may be primed for inflammatory cell death pathways.

We further noted enrichment of multiple components of the MHC-I antigen presentation pathway, including HLA-A/B/C, immunoproteasome subunits (PSMB/PSME), and the TAP1/2 peptide loading complex. MHC-II components were likewise increased (**Fig. 4C**). This pattern indicates enhanced antigen processing, transport, and presentation in the tumor relative to NAT, consistent with heightened interferon/hypoxia signaling and an inflammatory tumor milieu. Cell-type-oriented marker analysis revealed enrichment of macrophage (e.g., ARG1, CD14, CD163, TLR2), dendritic cell (e.g., CD209, ITGAX, SIRPA), and neutrophil (e.g., ELANE, MPO) markers in both tumor and PC relative to NAT, supporting active innate immunity in these compartments (**Fig. 4D**). Cytotoxic granule proteins (e.g., GZMA, GZMK, PRF1) were likewise elevated in the tumor and PC. Notably, T cell markers (e.g., CD3D, CD3E, CD4, CD8A) were more pronounced in the PC compared to both tumor and NAT, suggesting enriched T cell presentation and activity at the tumor boundary. Complement pathway proteins also showed increased abundance in the tumor and PC, consistent with innate immune activation in these regions.

Further, the tumor showed pronounced upregulation of key drivers of cell-cycle progression (**Fig. 5A**, blue), notably CCND1 (cyclin D1), cyclin-dependent kinases (CDK1/2), PLK1, and multiple minichromosome maintenance (MCM) complex members (MCM2/4/6/7), which support origin licensing and initiation of DNA replication.^40^ Concordantly, proliferating cell nuclear antigen (PCNA), a core replication factor, exhibited upregulation in the tumor. In parallel, angiogenesis-related factors were strongly elevated, consistent with the hallmark hypervascularity of ccRCC.^41^ Pro-angiogenic growth factors (e.g., VEGFA, TGFβ1), receptors (e.g., FLT1/4, MET), and members of the pro-angiogenic angiopoietin family (ANGPT2, ANGPTL2/4) were elevated in the tumor (**Fig. 5A**, red). Together, these data portray the tumor as both proliferative and strongly pro-angiogenic.

**Figure 5.**
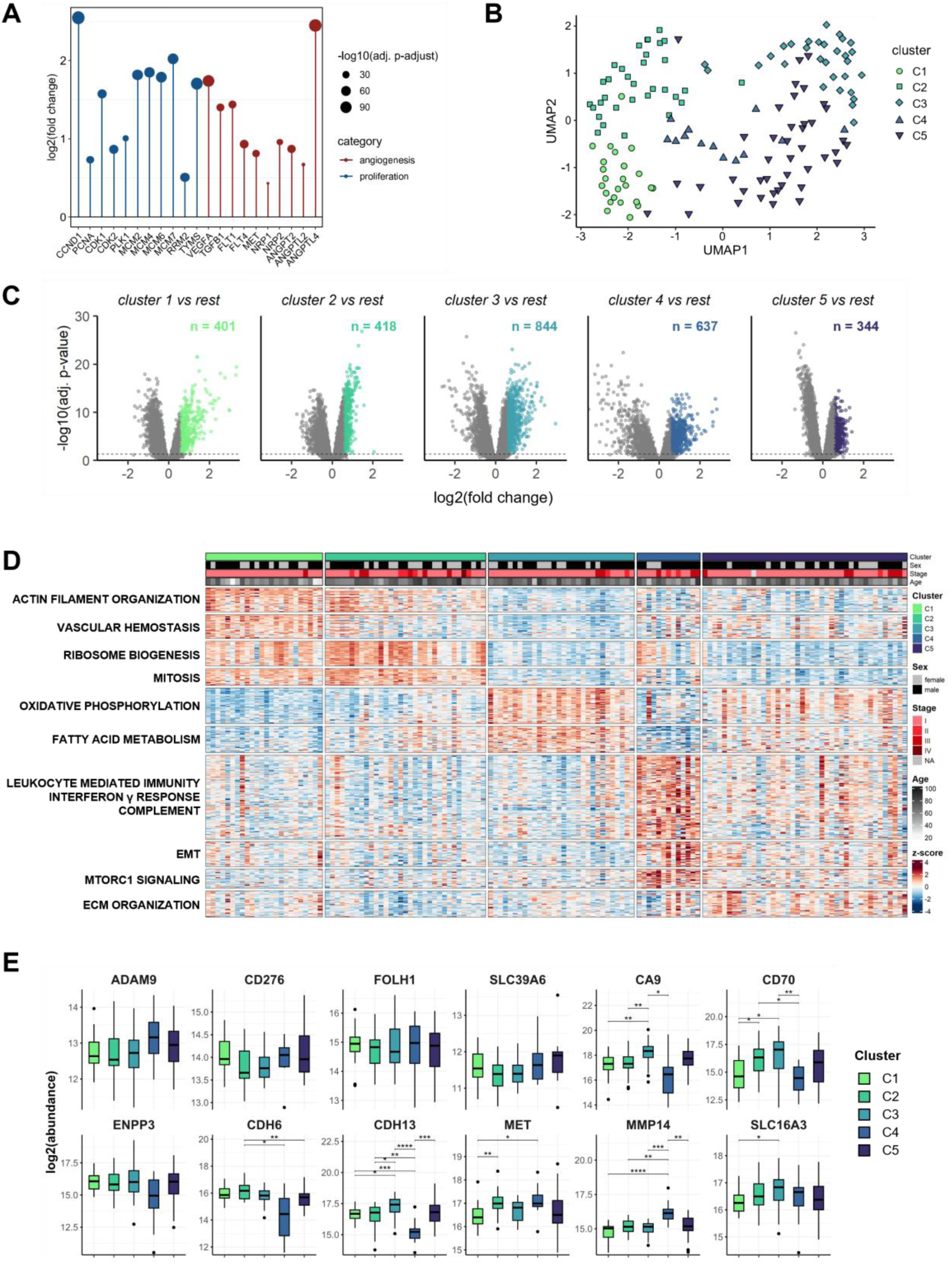
Sporadic ccRCC tumor clusters: **A** Dotplot of significantly upregulated proteins involved in angiogenesis (right) and proliferation (right) in the tumor (imputed dataset, fold change ≥ 1.5, BH adj. p-value ≤ 0.05). **B** Uniform Manifold Approximation and Projection (UMAP) of tumor samples (colors indicated the different tumor clusters). **C** Volcano plots showing differentially regulated proteins after limma analysis within the respective tumor clusters. Colored dots represent upregulated proteins (fold change ≥ 1.5, BH adj. p-value ≤ 0.05). **D** Heatmap of tumor samples showing enrichment of pathways in the different tumor clusters (for pathway analysis, Gene Ontology (GO), Hallmark gene sets (MSigDB), and Curated Cancer Cell Atlas (MSigDB) gene sets were use). **E** Boxplot showing the log2 abundance of selected ADC / theranostic target candidates in tumor clusters; *p < 0.05; **p < 0.01; ***p < 0.001; ****p < 0.0001 (Welch’s ANOVA, Games-Howell pairwise tests).

### Proteomic Subtyping Identifies Five Distinct Tumor Clusters in Sporadic ccRCC

To examine the intrinsic heterogeneity of sporadic ccRCC tumors beyond their primary tumor microenvironmental context, we performed an unsupervised proteomic subtyping analysis. Monte Carlo reference-based consensus clustering (M3C) identified five tumor clusters (C1– C5; **Fig. S4**). Uniform Manifold Approximation and Projection (UMAP; **Fig. 5B**) and PCA (**Fig. S5**) revealed a separation for C1–3 and C5, whereas C4 showed a more diffuse distribution, potentially indicating higher intra-group heterogeneity. Differential abundance analysis and pathway enrichment analyses uncovered distinct proteomic signatures across the five clusters (**Fig. 5C-D**). C1 was characterized by a strong proliferative and cellular organization signature, with enrichment in proteins associated with core processes like actin filament organization, ribosome organization, and mitosis, alongside an enrichment for venous hemostasis.

C2 shared much of the proliferative signature of C1 but was notably distinguished by a depletion in vascular hemostasis proteins.C3 represented a unique metabolic phenotype, exhibiting strong elevation in proteins linked to canonical energy production, specifically oxidative phosphorylation and fatty acid metabolism. C4 displayed a clear immune-invasive microenvironment signature, highly enriched in immune system processes, including leukocyte-mediated immunity, interferon-γ response, and complement system. The simultaneous elevation of proteins implicated in epithelial-mesenchymal transition (EMT) and mTORC1 signaling suggests an aggressive, invasive, and highly reactive TME phenotype for this group.^42,43^ C5 presented a more mixed or intermediate proteomic profile, with moderate enrichment of ECM proteins compared to the remaining clusters. These cluster patterns align with the proteomic subtypes reported in a recent proteogenomic study, which classified ccRCC into immune, metabolism, and ECM groups, corresponding broadly to our C4 (immune), C3 (metabolism), and C5 (ECM-rich) clusters.^44^ We next evaluated clinically relevant antibody-drug conjugate (ADC) targets currently investigated in ccRCC (**Fig. 5E**), and we observed cluster-dependent alterations in protein abundance. Notably, the immune-invasive C4, displayed reduced levels in CA9, CD70, CDH6, and CDH13, while MMP14 was elevated. This pattern suggests that C4 tumors may be less susceptible to CA9-, CD70-or cadherin-focused strategies, but could be candidates for MMP-targeting theranostic strategies.

### Distinct Proteolytic Processing in Different Tissues and Tumor Clusters

To capture endogenous proteolysis beyond fully tryptic peptides, we performed a semi-specific peptide search and quantified the resulting peptide species for each compartment. In total, we identified 19,701 semi-specific peptides, accounting for 20% of the total number of identified peptides (**Fig. 6A**). Cleavage motif analysis of significantly upregulated semi-specific peptides revealed similar motifs for both tumor (**Fig. 6B**) and PC (**Fig. 6C**). However, this did not allow for the assignment to a single protease or a group of proteases with confidence.

**Figure 6.**
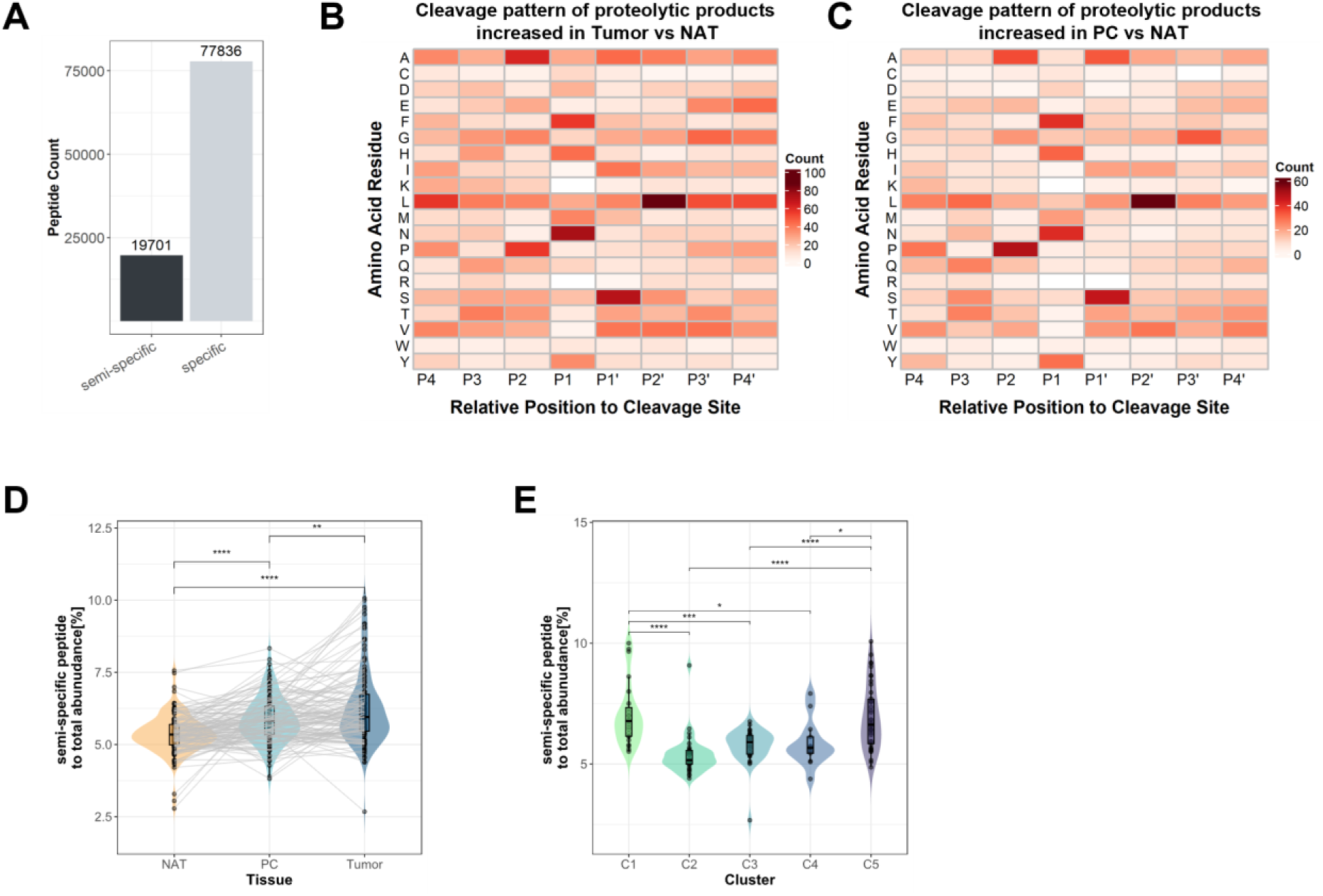
Semi-specific peptide analysis: **A** Barplot showing the number of identified semi-specific and specific peptides. **B, C** Heatmap showing proteolytic patterns of upregulated proteolytic products in tumor vs NAT (**B**) and PC vs NAT (**C**); fold change ≥ 2, BH adj. p-value ≤ 0.05. **D** Boxplot comparing the relative abundance of semi-specific peptides between tumor, PC, and NAT (grey lines connect samples from the same patient); **p < 0.01; ****p < 0.0001 (Dunn test, Tukey test). **E** Boxplot comparing the relative abundance of semi-specific peptides between tumor clusters C1–C5; *p < 0.05; ***p < 0.001, ****p < 0.0001 (Welch’s ANOVA, Games-Howell pairwise tests).

We next evaluated semi-specific peptide abundance by normalizing it to the total peptide abundance per sample. The proportional abundance of semi-specific peptides was significantly higher in the tumor compared to PC and NAT, consistent with our previous observations in syndromic ccRCC, pointing toward enhanced proteolytic processing in malignant tissue (**Fig. 6D**).^16^ Extending this analysis to tumor subtypes, we observed cluster-dependent differences. C1 and C5 exhibited the highest relative semi-specific peptide abundance, exceeding those of C2–C4 (**Fig. 6E**). Collectively, these results position proteolysis as a prominent and heterogeneous feature of sporadic ccRCC, suggesting elevated proteolysis within the tumor overall and subtype-specific variation.

## Conclusion

The present study constitutes the comprehensive proteomic profiling of 386 ccRCC FFPE tissue samples, comprising tumor, PC, and NAT. By achieving a deep proteome coverage of over 9,800 protein groups, a coherent view on sporadic ccRCC was delineated at both the tissue and the tumor-intrinsic level. The PC emerges as an ECM-rich, signaling-competent boundary enriched for scaffolding proteins, especially collagens, and it concentrates ECM-cell signaling molecules, including multiple integrins, LTBPs, and glypicans, consistent with enhanced growth factor sequestration and presentation at the tumor margin. In parallel, the PC exhibits signatures indicative of active remodeling. Semi-specific peptide profiling indicates heightened matrix turnover, and protease signatures evident with elevated MMPs (notably MMP7), balanced by increased TIMPs, suggesting spatially regulated proteolysis rather than unrestricted degradation. These features mirror those observed in our cohort of VHL disease-associated ccRCC, indicating a conserved boundary biology.^16^ The tumor proteome, in contrast, reflects the canonical pseudo-hypoxic metabolic state, with broad upregulation of glycolysis and hypoxia markers and depletion of aerobic metabolic pathways, including TCA cycle and electron transport chain components, alongside strong proliferative and pro-angiogenic signaling. Analysis of immune system-related processes points to heightened antigen-processing/presentation and inflammasome/pyroptosis signatures in tumors, while the PC concentrates T cell markers, consistent with an immune-active interface at the tumor margin. Mechanotransduction modules exhibit modest PC elevation, but their signature is dominated by inter-patient variance, implying that patient-level mechano-phenotypes potentially shape boundary remodeling beyond tissue identity. Unsupervised tumor subtyping resolves five tumor clusters spanning proliferative/mitotic (C1, C2), metabolic (C3), immune/EMT/mTORC1 (C4), and ECM-leaning (C5) programs. Semi-specific peptide abundance varies among clusters and shows highest abundance within the proliferative/metabolic C1 and the ECM-enriched C5. These molecular profiles align with the proteomic subtypes identified in a recently published large proteogenomic ccRCC study and bear practical implications, such as variable abundance of current ADC target candidates across clusters.^44^

Collectively, our compartment-resolved proteomic analysis of sporadic ccRCC reveals a conserved yet heterogeneous tumor boundary and highlights remodeling-and signaling-associated processes that may inform surgical strategy. Concurrently, identification of five discrete tumor clusters with divergent proteomic fingerprints establishes the foundation for subtype-aware therapeutic approaches.

## Data availability

The proteomic raw data will be available under restricted access for medical data protection reasons upon journal submission. Until then, please contact the corresponding author for access.

## Acknowledgments

We thank Katja Gräwe, Marlene Stigler, Bettina Wehrle and Hanna Kraatz for expert technical assistance. In addition, we would like to express our gratitude to all members of our laboratories for their helpful discussions and support.

## CRediT authorship contribution statement

**Tobias Feilen:** conceptualization, methodology, formal analysis, investigation, writing–original draft, writing–review and editing, visualization. **Grigor Andreev:** investigation, writing–review and editing. **Manuel Rogg:** investigation, writing–review and editing. **Miguel Cosenza-Contreras:** formal analysis, visualization, writing–review and editing. **Niko Pinter:** writing– review and editing. **Max Zirngibl:** data curation. **Antonia Elsässer:** data curation. **Pitter Huesgen:** resources, writing–review and editing **Markus Grabbert:** data curation, writing– review and editing. **Christoph Schell:** conceptualization, resources, writing–review and editing. **Oliver Schilling:** conceptualization, project administration, investigation, resources, supervision, funding acquisition, writing–original draft, writing–review and editing.

## Declaration of generative AI in scientific writing

During the preparation of this work the author(s) used ChatGPT (OpenAI) in order to refine the scientific writing process, finding synonyms and different phrasings. After using this tool/service, the author(s) reviewed and edited the content as needed and take(s) full responsibility for the content of the publication.

## Abbreviations

ABC: Ammonium bicarbonate
ACN: Acetonitrile
ADC: Antibody–drug conjugate
BCA: Bicinchoninic acid
CAA: 2-Chloroacetamide
CDK4/6: Cyclin-dependent kinase 4/6
CPTAC: Clinical Proteomic Tumor Analysis Consortium
DIA: Data-independent acquisition
HIAR: Heat-induced antigen retrieval
HPLC: High-pressure liquid chromatography
iRT: Indexed retention time
LC-MS/MS: Liquid-chromatography tandem mass-spectrometry
Limma: Linear models of microarray analysis
Lys-C: Lysyl endopeptidase
MET: Hepatocyte growth factor receptor
MHC: Major histocompatibility complex
NAT: Non-malignant adjacent tissue
PASEF: Parallel accumulation-serial fragmentation
PC: Pseudocapsule
PYCARD: Apoptosis-associated speck-like protein containing a CARD
S100A4: Fibroblast-specific protein 1
SDS: Sodium dodecyl sulfate
SP3: Single-pot solid-phase-enhanced sample-preparation
SPD: Samples per day
SUN1/2: SUN domain protein 1/2
TAZ: WW domain-containing transcription regulator protein 1
TCEP: Tris(2-carboxyethyl)phosphine hydrochloride
Tris: Tris(hydroxymethyl)aminomethane
YAP1: Transcriptional coactivator YAP1.

## References

1. Bukavina, L. et al. Epidemiology of Renal Cell Carcinoma: 2022 Update. Eur. Urol. 82, 529–542 (2022).

2. Shimizu, T. et al. Molecular mechanism of formation and destruction of a pseudo-capsule in clear cell renal cell carcinoma. Oncol. Lett. 27, 1–13 (2024).

3. Minervini, A. et al. Pathological characteristics and prognostic effect of peritumoral capsule penetration in renal cell carcinoma after tumor enucleation. Urol. Oncol. Semin. Orig. Investig. 32, 50.e15–50.e22 (2014).

4. Chen, J.-Y., Yiu, W.-H., Tang, P. M.-K. & Tang, S. C.-W. New insights into fibrotic signaling in renal cell carcinoma. Front. Cell Dev. Biol. 11, 1056964 (2023).

5. Wang, L. et al. Critical histologic appraisal of the pseudocapsule of small renal tumors. Virchows Arch. 467, 311–317 (2015).

6. Roquero, L., Kryvenko, O. N., Gupta, N. S. & Lee, M. W. Characterization of Fibromuscular Pseudocapsule in Renal Cell Carcinoma. Int. J. Surg. Pathol. 23, 359–363 (2015).

7. Xi, W. et al. Prognostic significance of pseudocapsule status in patients with metastatic renal cell carcinoma treated with tyrosine kinase inhibitors. Transl. Androl. Urol. 10, 4132141–4134141 (2021).

8. Kaelin, W. G. Molecular basis of the VHL hereditary cancer syndrome. Nat. Rev. Cancer 2, 673–682 (2002).

9. Haase, V. H. Renal cancer: Oxygen meets metabolism. Exp. Cell Res. 318, 1057–1067 (2012).

10. Hsieh, J. J. et al. Renal cell carcinoma. Nat. Rev. Dis. Primer 3, 17009 (2017).

11. Creighton, C. J. et al. Comprehensive molecular characterization of clear cell renal cell carcinoma. Nature 499, 43–49 (2013).

12. Choueiri, T. K. & Kaelin, W. G. Targeting the HIF2–VEGF axis in renal cell carcinoma. Nat. Med. 26, 1519–1530 (2020).

13. Higgins, D. F., Kimura, K., Iwano, M. & Haase, V. H. Hypoxia-inducible factor signaling in the development of tissue fibrosis. Cell Cycle Georget. Tex 7, 1128–1132 (2008).

14. Gilkes, D. M., Semenza, G. L. & Wirtz, D. Hypoxia and the extracellular matrix: drivers of tumour metastasis. Nat. Rev. Cancer 14, 430–439 (2014).

15. Cox, T. R. The matrix in cancer. Nat. Rev. Cancer 21, 217–238 (2021).

16. Feilen, T. et al. Proteomic characterization of the pseudocapsule of clear cell renal cell carcinoma in VHL disease reveals a distinct microenvironment at the tumor boundary zone. Neoplasia 68, 101214 (2025).

17. Hughes, C. S. et al. Single-pot, solid-phase-enhanced sample preparation for proteomics experiments. Nat. Protoc. 14, 68–85 (2019).

18. Skowronek, P. et al. Rapid and In-Depth Coverage of the (Phospho-)Proteome With Deep Libraries and Optimal Window Design for dia-PASEF. Mol. Cell. Proteomics 21, 100279 (2022).

19. Demichev, V., Messner, C. B., Vernardis, S. I., Lilley, K. S. & Ralser, M. DIA-NN: neural networks and interference correction enable deep proteome coverage in high throughput. Nat. Methods 17, 41–44 (2020).

20. Kistner, F., Grossmann, J. L., Sinn, L. R. & Demichev, V. QuantUMS: uncertainty minimisation enables confident quantification in proteomics. 2023.06.20.545604 Preprint at 10.1101/2023.06.20.545604 (2023).

21. Accurate Proteome-wide Label-free Quantification by Delayed Normalization and Maximal Peptide Ratio Extraction, Termed MaxLFQ* - Molecular & Cellular Proteomics. https://www.mcponline.org/article/S1535-9476(20)33310-7/fulltext.

22. Rohart, F., Gautier, B., Singh, A. & Cao, K.-A. L. mixOmics: An R package for ‘omics feature selection and multiple data integration. PLOS Comput. Biol. 13, e1005752 (2017).

23. Smyth, G. K. limma: Linear Models for Microarray Data. in Bioinformatics and Computational Biology Solutions Using R and Bioconductor (eds. Gentleman, R., Carey, V. J., Huber, W., Irizarry, R. A. & Dudoit, S.) 397–420 (Springer, New York, NY, 2005). doi:10.1007/0-387-29362-0_23.

24. Wu, T. et al. clusterProfiler 4.0: A universal enrichment tool for interpreting omics data. The innovation 2, (2021).

25. Ashburner, M. et al. Gene Ontology: tool for the unification of biology. Nat. Genet. 25, 25– 29 (2000).

26. Liberzon, A. et al. The Molecular Signatures Database Hallmark Gene Set Collection. Cell Syst. 1, 417–425 (2015).

27. Tyler, M. et al. The Curated Cancer Cell Atlas provides a comprehensive characterization of tumors at single-cell resolution. *Nat*. Cancer 6, 1088–1101 (2025).

28. Stekhoven, D. J. & Bühlmann, P. MissForest—non-parametric missing value imputation for mixed-type data. Bioinformatics 28, 112–118 (2012).

29. Petrov, P. B., Considine, J. M., Izzi, V. & Naba, A. Matrisome AnalyzeR – a suite of tools to annotate and quantify ECM molecules in big datasets across organisms. J. Cell Sci. 136, jcs261255 (2023).

30. Hoffman, G. E. & Schadt, E. E. variancePartition: interpreting drivers of variation in complex gene expression studies. BMC Bioinformatics 17, 483 (2016).

31. John, C. R. et al. M3C: Monte Carlo reference-based consensus clustering. Sci. Rep. 10, 1816 (2020).

32. Clark, D. J. et al. Integrated Proteogenomic Characterization of Clear Cell Renal Cell Carcinoma. Cell 179, 964–983.e31 (2019).

33. Voß, H. et al. HarmonizR enables data harmonization across independent proteomic datasets with appropriate handling of missing values. Nat. Commun. 13, 3523 (2022).

34. Andreev, G. et al. Spatial Correlation of the Extracellular Matrix to Immune Cell Phenotypes in the Tumor Boundary of Clear Cell Renal Cell Carcinoma Revealed by Cyclic Immunohistochemistry. Lab. Invest. 105, 104130 (2025).

35. Kalamajski, S. & Oldberg, Å. The role of small leucine-rich proteoglycans in collagen fibrillogenesis. Matrix Biol. 29, 248–253 (2010).

36. Robertson, I. B. et al. Latent TGF-β-binding proteins. Matrix Biol. 47, 44–53 (2015).

37. Filmus, J. Glypicans in growth control and cancer. Glycobiology 11, 19R–23R (2001).

38. Broz, P. & Dixit, V. M. Inflammasomes: mechanism of assembly, regulation and signalling. Nat. Rev. Immunol. 16, 407–420 (2016).

39. Tsuchiya, K. Switching from Apoptosis to Pyroptosis: Gasdermin-Elicited Inflammation and Antitumor Immunity. Int. J. Mol. Sci. 22, 426 (2021).

40. Blow, J. J. & Hodgson, B. Replication licensing — Origin licensing: defining the proliferative state? Trends Cell Biol. 12, 72–78 (2002).

41. Baldewijns, M. M. et al. High-grade clear cell renal cell carcinoma has a higher angiogenic activity than low-grade renal cell carcinoma based on histomorphological quantification and qRT–PCR mRNA expression profile. Br. J. Cancer 96, 1888–1895 (2007).

42. Kucejova, B. et al. Interplay Between pVHL and mTORC1 Pathways in Clear-Cell Renal Cell Carcinoma. Mol. Cancer Res. 9, 1255–1265 (2011).

43. Ganner, A. et al. VHL suppresses RAPTOR and inhibits mTORC1 signaling in clear cell renal cell carcinoma. Sci. Rep. 11, 14827 (2021).

44. Qu, Y. et al. A proteogenomic analysis of clear cell renal cell carcinoma in a Chinese population. Nat. Commun. 13, 2052 (2022).

