## Supplementary Tables and Figures for "Proteomic characterization of sporadic clear cell renal cell carcinoma reveals a matrix-dense pseudocapsule and heterogeneous tumor subtypes"

**Supplementary Table Legends**

| Variable | N | **Overall**, N = 153[1] | **female**, N = 58[1] | **male**, N = 95[1] | **p-value**[2] |
|---|---|---|---|---|---|
| **Age** | 153 | 64 (57, 72) | 66 (57, 77) | 63 (57, 70) | 0.050 |
| **Stage** | 153 | | | | 0.2 |
| I | | 123 (80%) | 50 (86%) | 73 (77%) | |
| II | | 10 (6.5%) | 1 (1.7%) | 9 (9.5%) | |
| III | | 18 (12%) | 7 (12%) | 11 (12%) | |
| IV | | 1 (0.7%) | 0 (0%) | 1 (1.1%) | |
| NA | | 1 (0.7%) | 0 (0%) | 1 (1.1%) | |
| **Side** | 153 | | | | 0.8 |
| left | | 63 (41%) | 23 (40%) | 40 (42%) | |
| right | | 90 (59%) | 35 (60%) | 55 (58%) | |
| **Tissue Samples** | 386 | | | | >0.9 |
| Tumor | | 142 (37%) | 51 (36%) | 91 (37%) | |
| NAT | | 121 (31%) | 46 (32%) | 75 (31%) | |
| PC | | 123 (32%) | 46 (32%) | 77 (32%) | |

[1]Median (IQR); n (%)

[2]Wilcoxon rank sum test; Fisher's exact test; Pearson's Chi-squared test

### Supplementary figures and legends

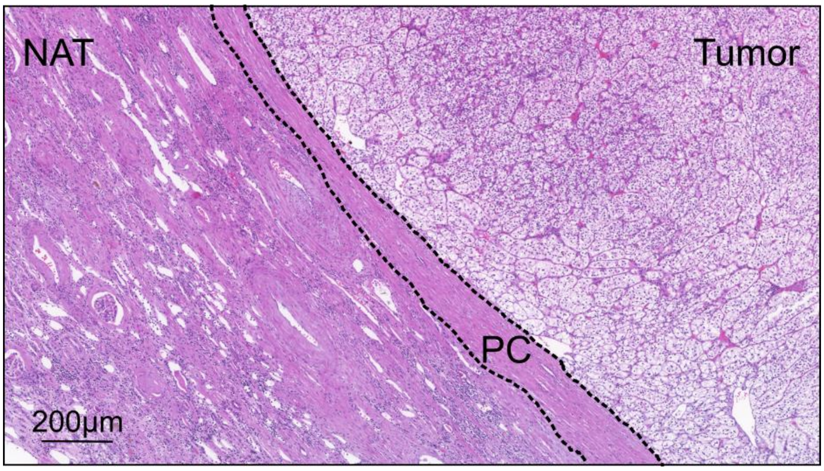

**Figure S1**. Representative H&E image of the ccRCC fibrous pseudocapsule (PC; dashed line), tumor, and NAT.

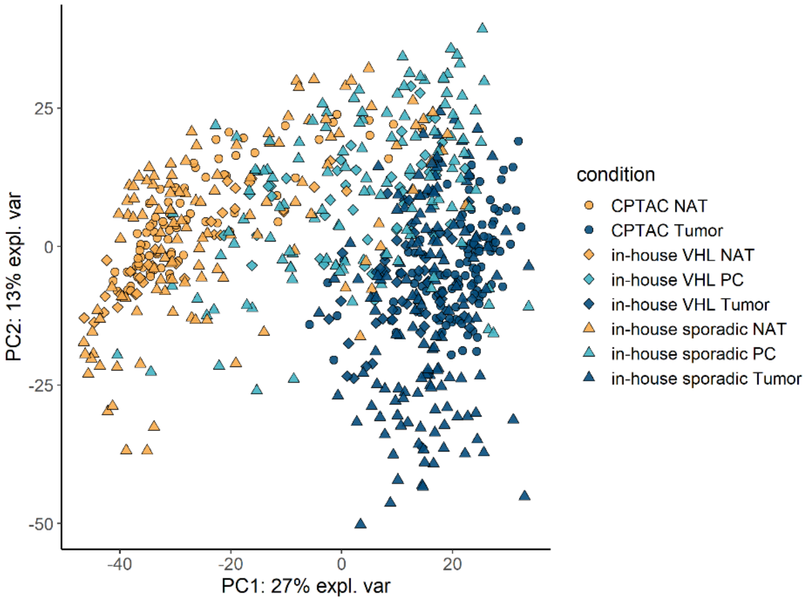

**Figure S2**. Principal component analysis (PCA) showing tumor, NAT, and PC samples of our in-house sporadic and syndromic ccRCC and the public CPTAC ccRCC dataset.

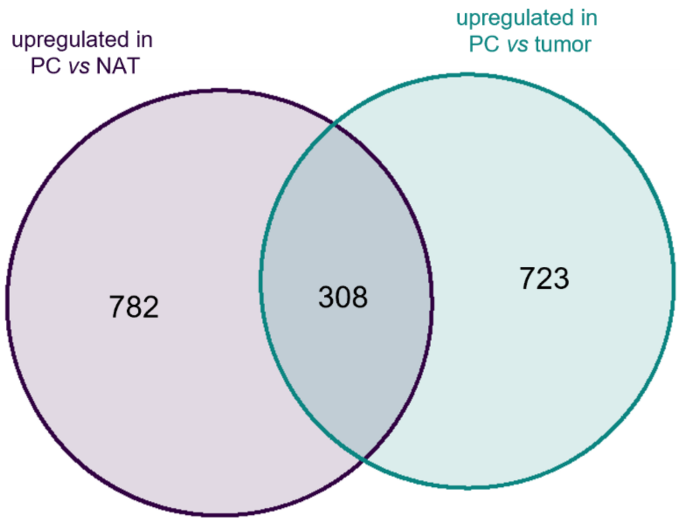

**Figure S3.** Venn diagram of upregulated proteins in PC vs NAT and PC vs tumor (imputed dataset, fold change > 1.5, BH adj. p-value < 0.05).

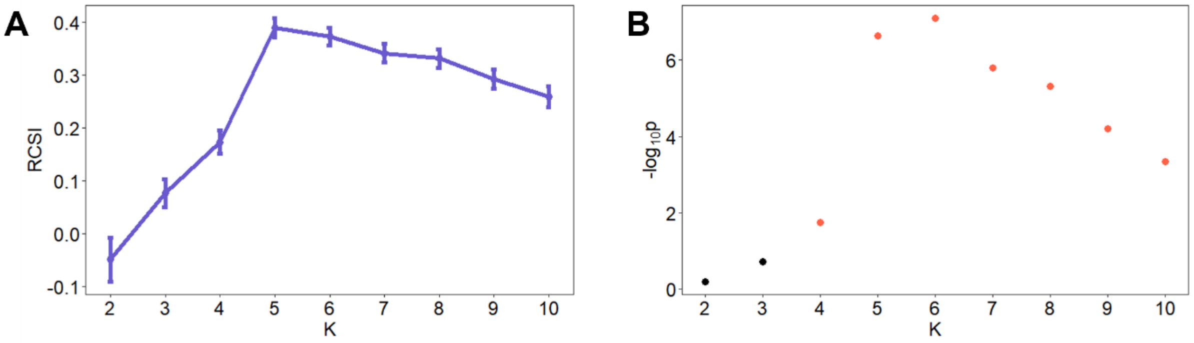

**Figure S4. M3C clustering results: A** RCSI (relative cluster stability index) indicates that the optimal number of clusters (K) is found for K=5. **B** M3C significance plot across K. Solutions become significant from K=4 (red points) and peak at K=6, supporting a meaningful cluster structure at K=5–6.

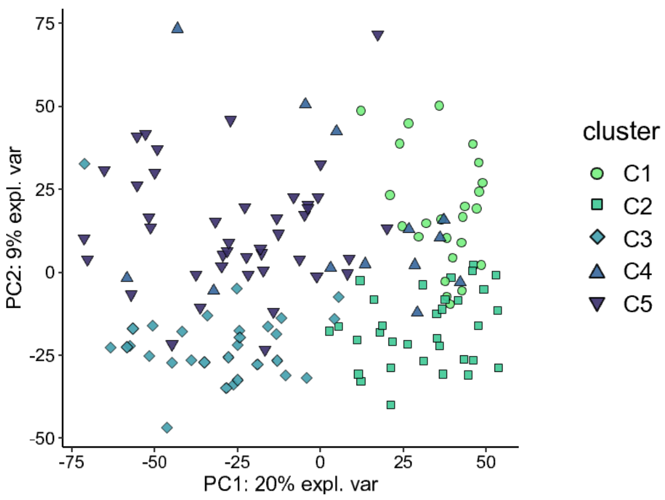

**Figure S5**. Principal component analysis (PCA) of tumor samples (colors indicated the different tumor clusters).
